# Visual neurons recognize complex image transformations

**DOI:** 10.1101/2024.06.10.598314

**Authors:** Masaki Hiramoto, Hollis T. Cline

## Abstract

Natural visual scenes are dominated by sequences of transforming images. Spatial visual information is thought to be processed by detection of elemental stimulus features which are recomposed into scenes. How image information is integrated over time is unclear. We explored visual information encoding in the optic tectum. Unbiased stimulus presentation shows that the majority of tectal neurons recognize image sequences. This is achieved by temporally dynamic response properties, which encode complex image transitions over several hundred milliseconds. Calcium imaging reveals that neurons that encode spatiotemporal image sequences fire in spike sequences that predict a logical diagram of spatiotemporal information processing. Furthermore, the temporal scale of visual information is tuned by experience. This study indicates how neurons recognize dynamic visual scenes that transform over time.

**One-Sentence Summary:** Complex visual scenes are encoded by plastic trajectories of spatiotemporal image sequences.

---

Natural visual scenes consist of sequences of images that transform in a complex manner over time. Dynamic temporal components of visual inputs are essential for object recognition, suggesting the circuitry underlying this function is robust. Studies in the visual system have suggested that a basic principle of sensory processing is that individual geometric components of complex visual stimuli are isolated and are then integrated to recompose scenes (*1–4*). On the other hand, how the temporal sequence of dynamic visual information is represented is poorly understood. To address this question, we analyzed receptive fields of optic tectal neurons. The optic tectum/superior colliculus is an evolutionarily conserved visual processing center in vertebrates(*5*), which plays important roles in attention (*6–8*), decision making (*9*), and prey capture (*10*). It is noteworthy that each of these functions requires processing temporal information (*11, 12*). Receptive field maps are known to encode different spatial features in the environment, such as ON/OFF stimuli or more complex features such as direction (*13, 14*). Here, we used unbiased sparse noise stimuli and reverse correlation to search stimulus space over extended response times. Our study revealed that tectal neurons recognize a variety of image sequences that transform over time, reflecting the temporal relationship of spike timing across neurons in the circuit. Furthermore, the encoded temporal dynamics are plastic and can be tuned to prevalent temporal dynamics in the environment. The data support a network model of how dynamic spatiotemporal information characteristic of natural scenes may be encoded over time.

## Results

### Temporal dynamics of receptive fields

Visual response properties were recorded from optic tecal neurons using juxtacellular multichannel recordings (Fig. S1), allowing us to record 2-4 units over extended periods (*15*). We first mapped the spatiotemporal receptive field properties of visual responses using classical methods, specifically, with cell-attached recordings to detect responses to small white square stimuli presented in a matrix of pseudorandom locations in the visual field (*16*) (Fig. S2A). The spatial receptive field (RF), shown as a pseudocolored map, represents neuronal firing in response to stimuli in a particular location (Fig. S2B). Spike numbers change over time, suggesting temporal dynamics of the spatial RFs, which differed depending on the locations of the stimulus within the visual field (Fig. S2A-ii,iii, S2B). The similarity of the spatial RFs rapidly decreased with longer time intervals between maps (Fig. S2C, D), also suggesting rapid temporal changes in the spatial RFs. Direction selective responses also displayed temporal dynamics in the preferred direction (Fig. S3A-C). Although these data suggest that individual optic tectal neurons may recognize temporally dynamic components of visual stimuli, the spatiotemporal dynamics of the maps varied over across three repetitions of the same stimuli (Fig. S2B). We therefore improved the strategy to assess the spatiotemporal dynamics of RF maps. We used an unbiased sparse noise visual stimulation protocol, presenting a random series of stimuli composed of two squares in different configurations(*3*) and averaged responses to test stimuli over a large number of spatial and temporal combinatorial settings. Images of stimuli were interleaved with 400-1000ms intervals to detect extended temporal response dynamics (Fig. 1A). RF maps were calculated by reverse correlation (*3, 17*) and presented as a spike triggered average (STA) where the image in each time window is the average of stimuli that evoked spikes at 0 ms (*18*). We found that RF maps changed significantly over time (Fig. 1B). The optimal stimulus identified by reverse correlation typically began with an ON stimulus followed by an OFF stimulus which dominated during the last 100-150 ms of the stimulus. In addition to the transition from ON to OFF responses, we observed multipolar spatiotemporal RF dynamics during the transition period, including dipole responses of transitions from ON to ON/OFF to OFF and tripole responses of ON to ON-OFF-ON to OFF or ON to OFF-ON-OFF to OFF responses (Fig. 1B, top panel: two, three). Another feature of these dynamic RFs is the motion of the RF map components, assessed as changes in the gravity center of the ON or OFF responses. For instance, in one set of RFs, ON and OFF components both shifted (Fig. 1B, bottom panel: shift), while in a second set, the vector of the ON-OFF line rotated (Fig. 1B, bottom panel: rotate). In a third set, the pattern transformed in a complex manner (Fig. 1B, bottom panel: complex). Half of the responses are dipoles, 4% are tripoles and 15% are more complex (Fig. 1C, left). Furthermore, of the dynamic RFs, 30% showed no change in the gravity center, while 70% either shifted, rotated or showed more complex types of motion (Fig. 1C right: stable, shift, rotate, complex respectively). Together these data indicate that optic tectal neurons recognize complex image sequences in the temporal dimension.

**Fig. 1.**
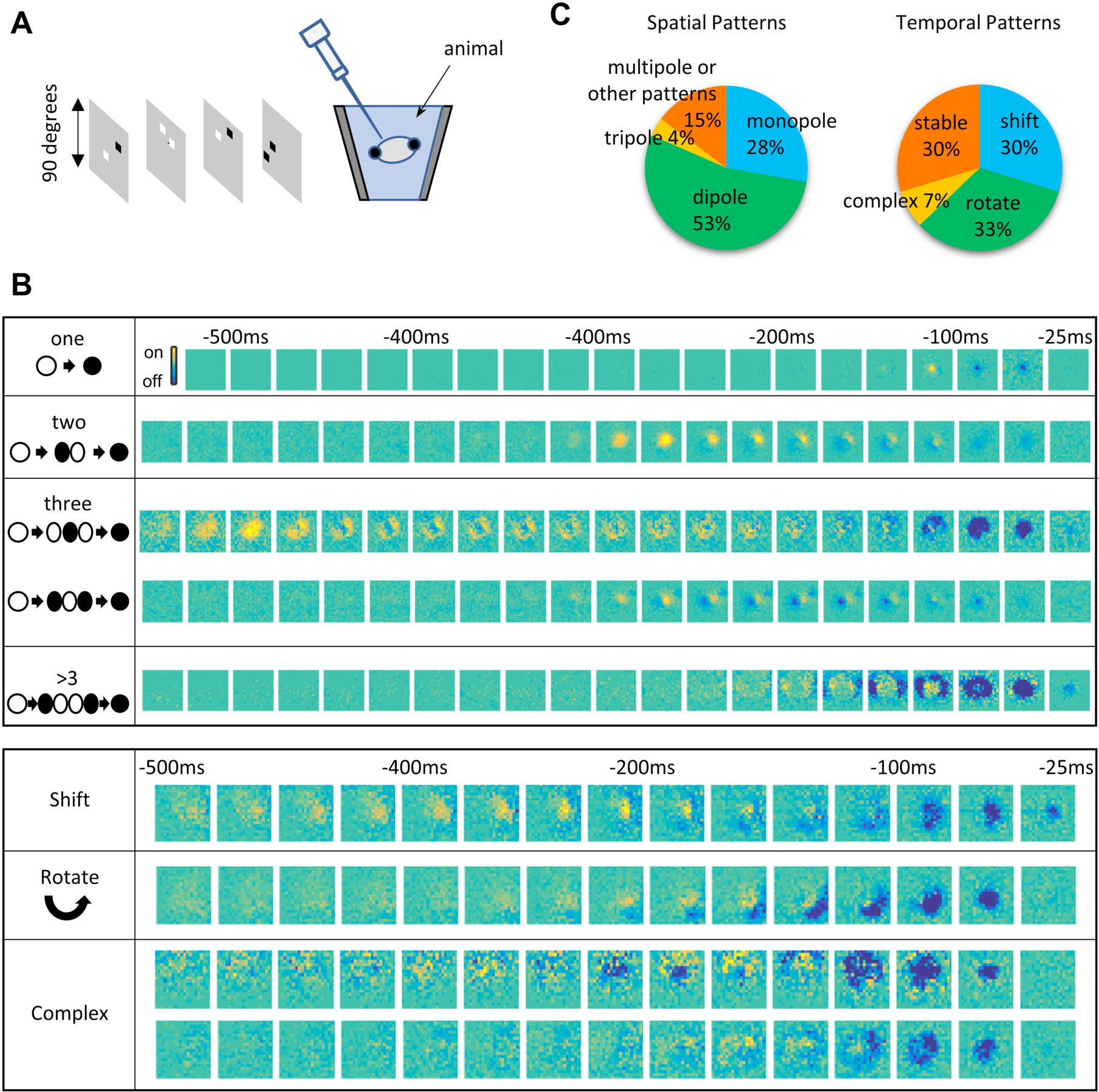
Tectal cell visual response properties change over time. **A.** Recording setup. Animals were placed on a stage in the center of a recording chamber equipped with a rear-projector screen on the side of the chamber. Black and white squares represent stimuli used to identify tectal cell response properties. Neuronal activity was recorded with juxtacellular glass electrodes in response to an unbiased sparse noise visual stimulation protocol, consisting of 4000-16000 randomly generated images of combinations of two white or black squares on a gray background, presented for 400-600ms without a gap. **B.** Spatiotemporal response properties change over time, displayed as a Spike Triggered Average (STA). **(top)** Polar spatiotemporal responses, showing increasing complexity of optimal stimuli from one, two, three or more poles. **(bottom)** Categories of temporal response dynamics. Shift: parallel shift without changing the vector of ON to OFF poles. Rotate: two examples in which the vector connecting ON and OFF poles spins. Complex: other complex changes, such as changes in distributed ON and OFF responses over time. **C.** Percentages of spatial patterns (right) and temporal patterns (left).

### Determining spatiotemporal filters

Recognition of complex motion stimuli, such as rotation, may employ specific spatiotemporal filters. Although several computational models have been proposed for a circuit-based mechanism to recognize rotational stimuli (*19, 20*), and specifically spinning stimuli, made famous by the perception of wagon wheel aliasing (*21, 22*), neural mechanisms that recognize rotational motion have not been described. Our unbiased stimulation protocol indicated that a significant proportion of neurons encode rotation in either direction (Fig. 1B, C (rotate), 2A). We then tested whether neurons display a preference for rotational direction by presenting a clockwise or counterclockwise rotating stimulus over several rotation speeds (Fig. 2B). Many units exhibited a preference for a rotation direction for at least one rotation speed (Fig. 2C-E, S4A). The experimentally observed rotation direction preference in response to the rotating stimulus was similar to the simulation generated from the cross-correlation between the optimal stimulus and the presented stimulus (Fig. S4B), validating the identification of the optimal stimulus. It is interesting that tectal neurons exhibited speed dependent changes in the preferred direction of rotational stimuli (Fig. 2C, right), similar to the perception of wagon wheel aliasing(*21, 22*). To explore the logical mechanism underlying rotation direction selectivity, we asked whether this rotation direction selectivity can be achieved by the summation of a series of linear direction selective responses. We presented drifting gratings moving clockwise (CW) or counterclockwise (CCW) positioned at 30° intervals to the animals recorded in Fig. 2C left (Fig. 2F). We compared the total number of spikes evoked by the two sets of stimuli moving in opposite directions (Fig. 2G, magenta and blue). The magenta and the blue lines overlapped. This indicates that the sum of the responses evoked by the series of individual stimuli was similar for both directions, showing no directional preference for the set of linear movements, in contrast to the robust response to the rotational stimulus (Fig. 2C, left). Population analysis showed that more neurons were selective for rotational direction rather than the set of linear movements (Fig. S4C). We detected no correlation between the rotational and linear direction selectivity with respect to speed dependent responses (Fig S4D, E). These results indicate that rotation direction selectivity is due to the integration of the sequence of image stimuli moving in a nonlinear spatial pattern and not the sum of linear direction-selective responses. Together these data provide evidence for the presence of spatiotemporal filters that recognize non-linear movements, through the detection of sequential changes in both spatial and temporal dimensions.

**Fig. 2.**
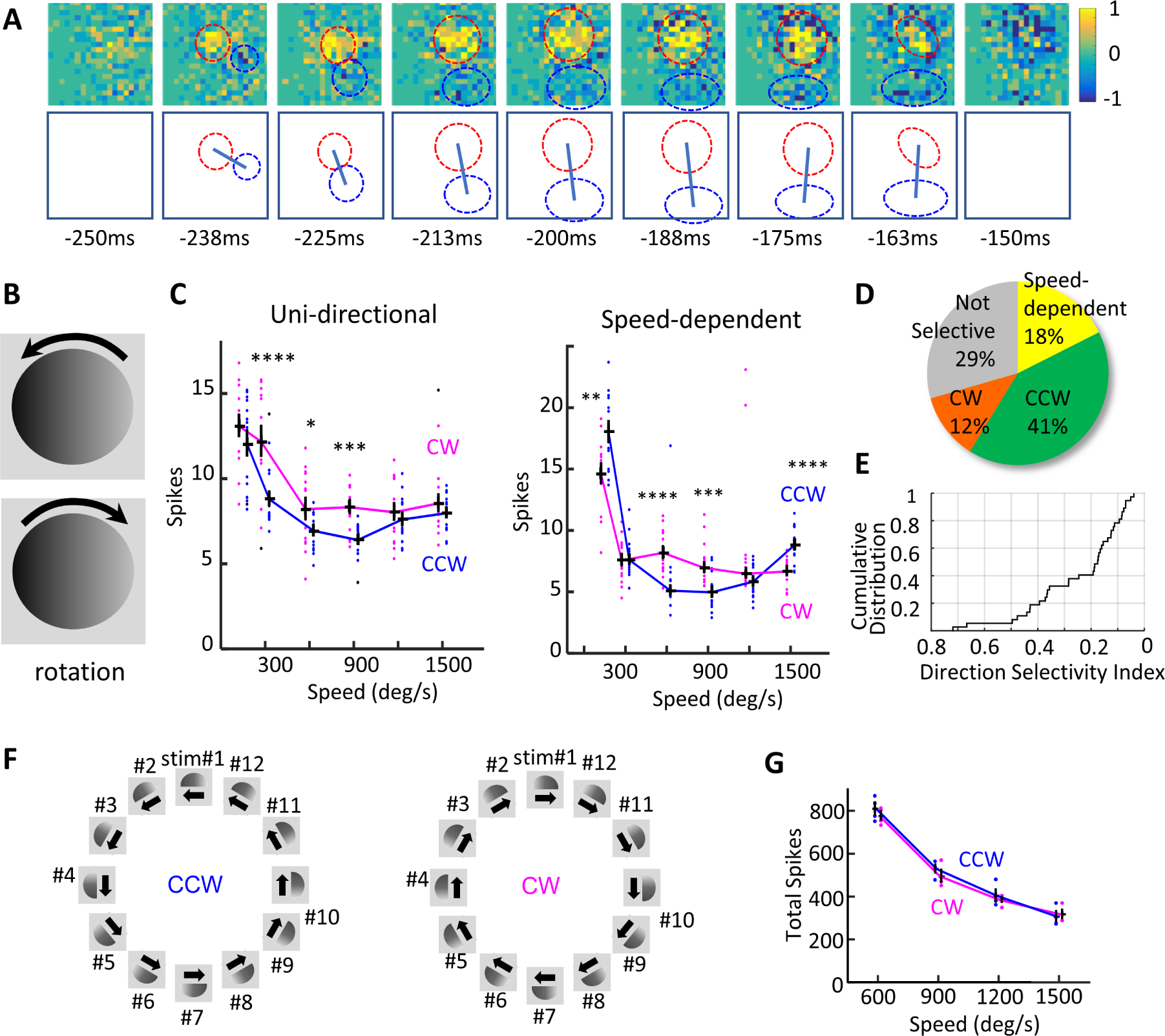
Tectal neurons detect direction selective rotating stimuli. **A.** Unbiased stimulus presentation identifies rotation as an optimal stimulus. Top: RF maps determined by reverse correlation analysis. Bottom: The axis connecting ON-OFF RF centers (red and blue, respectively) rotated over time. bin: 25ms. Values below RF maps are the ending timepoints of the 25 ms bins. Color scale: spike numbers normalized to the absolute maximum value. **B.** Stimuli used to test direction selectivity of the rotation response. A counterclockwise (CCW, top) or clockwise (CW, bottom) rotating circular gradient was presented in the RF. **C.** Number of spikes evoked by the rotating image at different speeds from two representative neurons shows directional selectivity for rotation. **(left)** An example of preference for CW rotation (magenta) compared to CCW (blue). **(right)** A neuron in which the preferred rotation direction showed speed-dependent switching between CCW (blue) and CW (magenta) rotation. Responses to CW and CCW rotation were significantly different at several rotation speeds. * p< 0.05, *** p< 0.001 bootstrap test, 10^5^ repeats, n=6 sessions. **D.** Percentages of neurons showing significant rotation direction selectivity (n=15 units). Neurons with P[#Spikes(CW)=#Spikes(CCW)]<0.05 and |DSI|≤0.1 at any rotation speed are counted as rotation direction selective. **E.** Cumulative distribution of rotational direction selectivity index at the optimal speed. (n=38 units). **F-G.** Rotation direction selectivity is not achieved by the summation of responses to a series of linear direction selective stimuli. **F**. Sets of stimuli to evaluate summation of linear direction selective responses. Drifting gratings were presented in semicircles of 12 orientations in the cell’s RF in CCW (blue) and CW (magenta) directions. **G.** Total number of spikes evoked by the sets of linear CW or CCW motion stimuli. DSI = 0.020 (600deg/s), 0.033 (900deg/s), 0.025(1200deg/s), -0.018 (1500deg/s). Compare with responses of the same cell to rotational stimuli (**C, left)**.

The experiments above suggest that rotation responsive neurons prefer a specific sequence of image stimuli. To test this directly we first recorded responses to an unbiased series of stimuli and determined the optimal stimulus sequence for the RF by reverse correlation (Fig. 3A). We then generated movies from the optimal, reverse or shuffled image sequences (Fig 3B-i, Fig. S5A) and recorded responses to the movies, plotted as the numbers of spikes binned in 100ms windows over the 600 ms movie (Fig. 3B-ii). Images presented in the optimal sequence evoked the largest number of spikes (blue), whereas the reverse sequence evoked the fewest spikes, and the shuffled image sequences evoked intermediate responses (Fig. 3B-ii,iii, Fig. S5A, B). Changes in radiance do not account for changes in responses to forward, reverse or shuffled stimulus sequences (Fig. S5C). Together, these data suggest that the neurons preferentially respond to the optimal sequence, discovered by unbiased stimulus presentation, but respond poorly to the reverse stimulus sequence. We evaluated whether neurons exhibit a preference for the image presentation sequence by defining an Image Sequence Selectivity Index (ISSI), [spikes(original) – spikes(reverse)]/[spikes(original) + spikes(reverse)], analogous to the spatial direction selectivity index. This analysis indicated that neurons showed a significant preference for the original image sequence compared to the reverse image sequence (Fig. 3C: CTL).

**Fig. 3.**
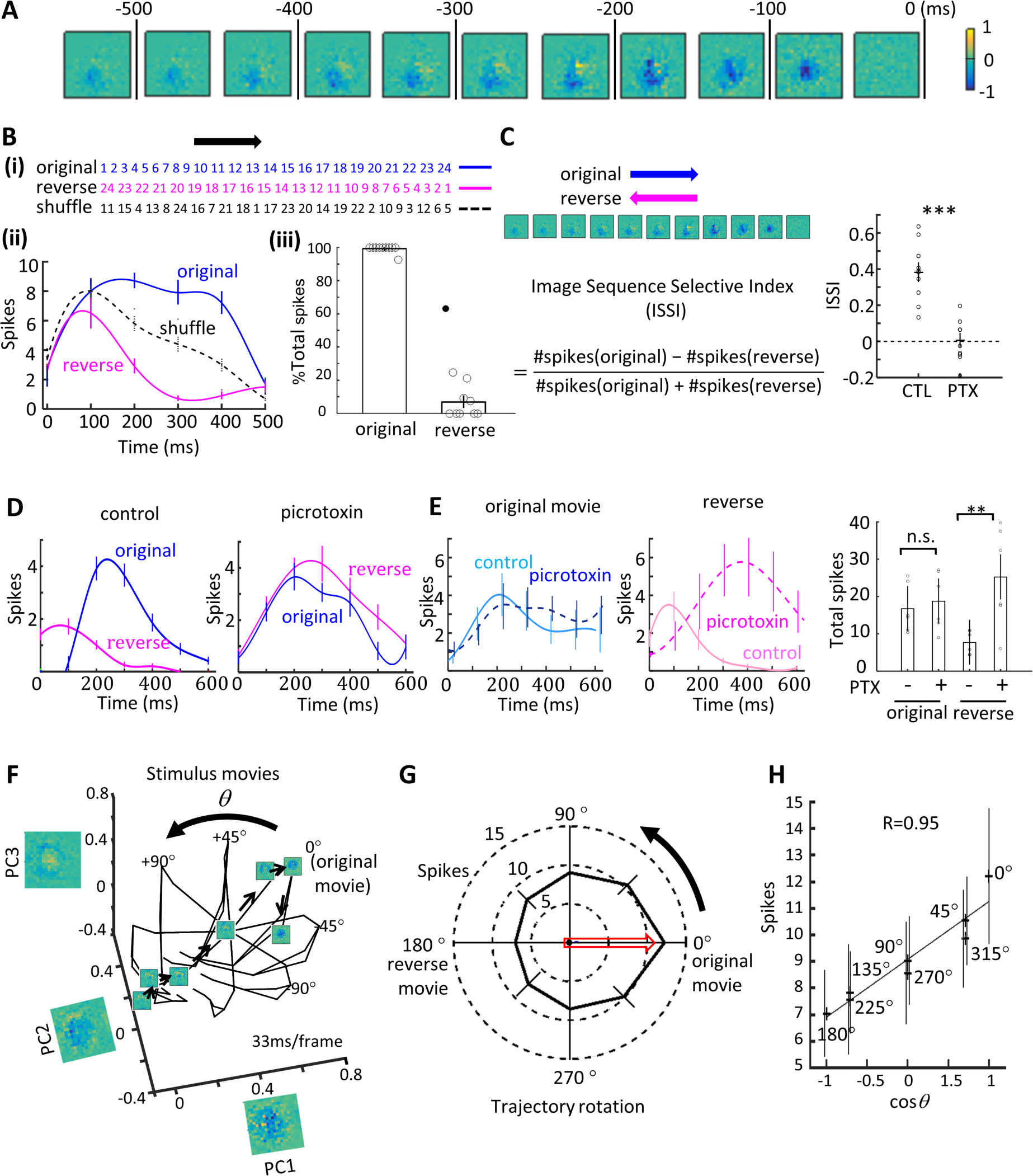
Tectal neurons recognize sequence-specific image transitions. **A**. RF maps of visually-evoked response dynamics over time determined by reverse correlation analysis. ON and OFF responses are shown in yellow and blue, respectively. The image sequence that evoked this response is termed the “original” sequence. **B.** Analysis of stimulus sequence specificity of visual response properties. **(i)** Sequence of frame numbers for the original and reverse stimulus sequences and a representative sequence of shuffled stimuli (blue, magenta and black respectively). **(ii)** Temporal profiles of responses to original, reverse and shuffled stimulus sequences (blue, magenta and black respectively) for a representative neuron. Spike numbers are pooled over 100ms intervals for the 600ms stimulation period. Mean ± SEM, n=10 stimulus sessions. See **Fig. S5A, B** for data pooled from N=10 neurons. **(iii)** Relative spike numbers evoked by original or reverse image sequences. For each unit, the maximum and minimum numbers of spikes evoked by the set of movies was normalized to 100 and 0%, respectively. The movie of the original image sequence consistently evoked the largest responses throughout the movie. Original: 99±0.77% total spikes, Reverse: 6.9±3.4% total spikes, Mean±SEM, N=10 neurons, including one outlier (filled circle, outside the interquartile range). **C.** Image Sequence Selectivity Index (ISSI), defined as [spikes(original) – spikes(reverse)]/[spikes(original) + spikes(reverse)]. Plots of ISSI from control and picrotoxin-treated animals. ISSI(control) = 0.38±0.05 (Mean±SEM, N=10 neurons), ISSI(PTX) = 0.0061±0.0378 (Mean±SEM, N= 8 neurons), p< 0.001. Paired t-test. **D.** Representative examples of temporal response profiles recorded from control (left) and 100µM picrotoxin-treated (right) animals for optimal and reverse stimulus sequences. Spike numbers are pooled over 100ms intervals for the 600ms stimulation period. Mean±SEM, N=20 stimulus sessions. **E.** Paired comparison of the response profiles in control and 100µM picrotoxin conditions (6 pairs) for optimal (left) and reverse (middle) movies sequences. Total number of spikes (right) for optimal and reverse movie sequences recorded in control or PTX conditions. ** p< 0.01. Paired t-test. n.s.= not significantly different. **F.** Transformation of the original image sequence into a trajectory of principal coordinates (axes with image frames representing the principal coordinates) PC1:40.9%, PC2:13.9%, PC3: 6.6%. A series of test movie stimuli was generated by rotating points in this trajectory along an axis, which was defined so that a 180-degree rotation generates a reverse image stimulation sequence. **G.** Polar plot of evoked spike numbers (Mean±SEM, N=6 neurons) in response to different trajectory rotations. The original movie (0° rotation) generated the most spikes. The reverse movie (180° rotation) generated the fewest spikes. **H.** Spike numbers encode rotation angle of the image sequence trajectory in principal coordinates. Plot shows a linear relationship between the cosine of rotation angles and total number of spikes (Mean±SD, R=0.95, P<0.001. ANOVA test).

The extended period of sustained spiking in response to the optimal image sequence compared to the reverse sequence suggests that mechanisms, such as inhibition, may be involved in generating the complex spatiotemporal response properties. We recorded responses to the optimal and reverse image sequences under control conditions and in the presence of picrotoxin (PTX, 100µM) to block GABAergic inhibition. PTX increased the response to the reverse movie sequence so that it is similar to the response to the optimal movie sequence (Fig. 3D, E). PTX significantly decreased the Image Sequence Selectivity Index compared to control (Fig. 3C), indicating a role for inhibition in the image sequence selectivity encoded in the temporally and spatially dynamic RFs.

Next, we investigated how the information space is represented by neural activity. To visualize transitions of spatiotemporal RF maps across time, we plotted the sequence of RF map transitions according to changes in their principal components over time (Fig. S5D). Since RF maps start from ON responses and end with OFF responses in most cells (Fig. 1B), the beginning and the end of the trajectories of many neurons converged (Fig. S5D red and blue segments, respectively). The locations of the intermediate segments were variable, reflecting the variation between RF maps, while some trajectories were shared by subsets of neurons (Fig. S5D arrowheads), suggesting common spatiotemporal stimulus features in the optimal stimuli across neurons. Together, this analysis, using unbiased stimulus presentation together with recording periods extending over hundreds of milliseconds, indicated that RF responses can encode complex spatiotemporal visual information.

To further explore how spatiotemporal image transformations are encoded by neurons, we generated principal coordinates from the RF maps in response the unbiased stimulus presentation and analyzed the responses to image sequences represented in the coordinates. We converted the original image sequence into a trajectory in the principal coordinate (Fig. 3F) and then modified the trajectory to generate a series of image sequences to present as test stimuli. First, we tested parallel shifts of the trajectory in each coordinate axis, which essentially corresponds to addition of a background image to the series of the images. As expected, parallel shifts of the trajectory did not significantly change the responses (Fig. S5E), likely because differences in luminance between image frames (dIM_n_ = IM_n+1_ – IM_n_) were unchanged(*23*). We next rotated the original trajectory to obtain a set of trajectories in which the angle is altered systematically (Fig. 3F), generating a second set of test image sequences. The response evoked by the original stimulus (Fig. 3G, 0 degrees) was larger than the responses to the test stimuli of rotated movies (Fig. 3G, 45-315deg). Again, movies of the original and reverse (180° rotation) stimuli evoked the greatest or smallest number of spikes, respectively, in the series of the stimuli. Furthermore, the numbers of spikes matched a linear function of cos(θ) (Fig. 3H), showing that neurons do not only recognize specific image sequences but are capable of encoding a series of movie stimuli following a linear function.

### Network activity underlying spatiotemporal filters

Next, we determined whether tectal neurons show sequences of activity consistent with encoded spatiotemporal information. We explored the neuronal activity patterns, detected by in vivo calcium imaging, in response to unbiased presentation of visual stimuli, as described above (Fig. 4A, B, S6A, B). This protocol allows us to identify preferred stimulus sequences for each neuron and determine the relative spike timing across neurons. We analyzed spikes from all neurons in a pairwise fashion to identify pairs of neurons with a consistent firing sequence (Fig. S6C-F). We found 28.7% of pairs of cells fire in a consistent sequential order. This was not due to differences in response time of the different neurons to visual stimuli because only 0.22% of pairs showed significant differences in the latency from visual stimulus presentation to response times between the neurons in the pair (Fig. S6C-F). It is likely that our analysis underestimated the population of neurons that fire in sequence because we image a fraction of the neurons in the tectal circuit. Pairs of neurons with a consistent firing sequence often formed trains, in which multiple neurons fire in sequence (Fig. 4C, magenta). This sequential firing pattern rarely emerges in rasters of shuffled data (Fig. S6G-I). The number of trains of sequentially active neurons was far in excess of that seen in the shuffled data (Fig. S6G, H).

**Fig. 4.**
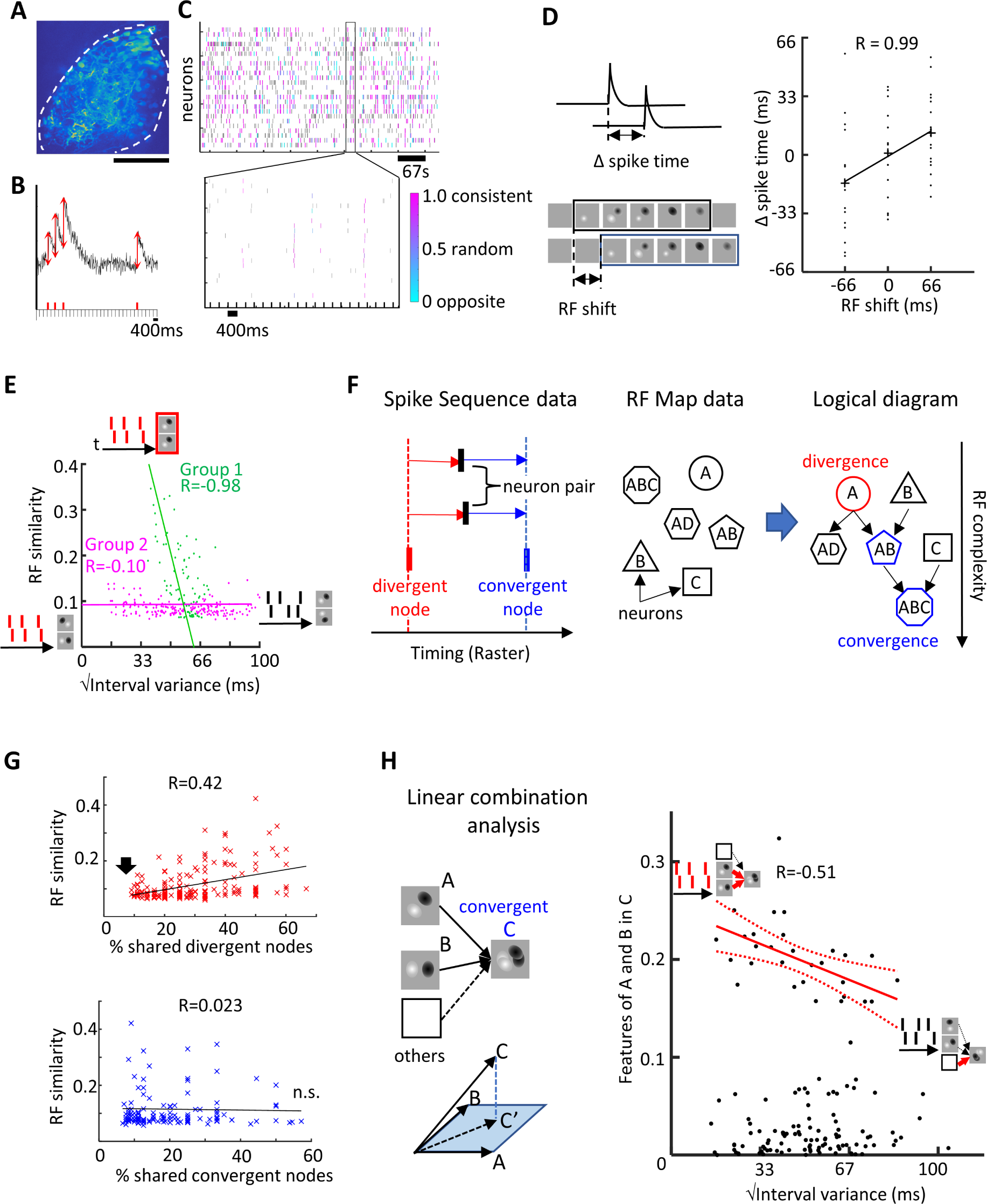
Multiunit spike sequence predicts a logical circuit diagram consistent with spatiotemporal RF map transformation. **A.** 2-photon image of GCaMP6f expression in the optic tectum. Dotted line: outline of the optic tectum. Scale bar: 200μm. **B.** Example of Ca^++^ spikes in response to unbiased stimulus presentation. Ticks on the X axis show timing of stimuli that generated Ca^++^ spikes. See **Fig. S6A, B** and Methods for spike detection criteria. **C.** Raster plot of Ca^++^ spikes. Color coded rasters show spikes from cells that fire in a reproducible sequence with other cells within a 100ms time window. **D. (left, top)** Schematic of the representative relationship between sequential spiking activity in neurons and **(left, bottom)** change in RF properties determined by reverse correlation of unbiased stimulus presentation over a similar time frame. **(right)** Plot of the temporal shift of the RF, determined by 3-dimensional cross-correlation, and the mean difference in spike timing between all pairs of neurons in a representative animal (see **Fig. S6J** for pooled data from N=4 animals.) **E.** Plot of RF similarity (cross-correlation of RF maps) versus square root of the variance in spike timing intervals. Each dot represents a pair of cells from an animal (see **Fig. S6L** for pooled data). Insets show schematics of spike intervals and RF maps. Red ‘spikes’ have consistent spike intervals, black ‘spikes’ have varying spike intervals. RF maps boxed in red are similar. Unboxed RF maps are dissimilar. **F.** Logical diagram inferred from calcium imaging data on spike sequence and RF maps. **(left)** Use of spike sequence and temporal coherence data to predict information flow between neurons. When a neuron (red or blue raster) consistently fires earlier or later than a pair of neurons (black raster), this neuron is modeled as an earlier, divergent or a later, convergent node. (**middle**) RF features predict information flow across neurons. RF features, depicted as letters, in different neurons, represented by different shapes. **(right)** Logical diagram modeling increased RF complexity, generated using information about spike sequence and RF features of neurons, which predict divergent and convergent nodes. **G.** The proportion of shared divergent nodes correlates with RF similarities in pairs of neurons (R=0.42) **(top)**; the proportion of shared convergent nodes does not correlate with RF similarity (R=0.023) **(bottom)**. N=52 cells from three animals. **H. (left)** Linear combination analysis of the RF map data. The optimal linear combination of image A and image B for the representation of image C was obtained as C’ by Moore-Penrose inverse matrix (sA + tB + X= C). **(right)** Plot of features of RF maps, A and B, present in map C, determined as |C’|^2^ / |C |^2^ from the linear combination analysis, versus square root of the variance of the interval between spiking. Data values from Y>0.1 were used for the correlation analysis. N=52 cells from three animals.

If the firing sequence of neurons reflects the timeline of visual information, the temporal relationship of spikes across neurons would be expected to predict the logical relationship of the information encoded by the neurons. Previous studies suggested the presence of neurons that fire in a reproducible temporal sequence (*24, 25*) and modeling studies predicted the potential of this type of network to process temporal information (*12*).We tested whether pairs of neurons with consistent intervals in spike timing show corresponding temporal dynamics in their RFs, as schematized in Fig. 4D, left. We found that the sequence of firing across neurons was correlated with the temporal dynamics of the spatiotemporal RFs calculated by three-dimensional cross-correlation (Fig. 4D, right, Fig. S6J), suggesting that visual information is transferred between neurons. To pursue the correspondence between relative spike timing and temporal dynamics of the RF maps, we plotted the relationship between the similarity of the spatiotemporal RF patterns and the consistency of sequential spike activity for each pair of neurons, quantified as the square root of the variance of the difference in spike intervals between two neurons. Notably, two clusters emerged in the plot (Fig. 4E, S6K, L). In one cluster of neuron pairs (Fig. 4E magenta), the RF maps showed no similarity even though some pairs are highly synchronized. In the other cluster (Fig. 4E green), the similarity of RFs between the neurons and the temporal consistency of their spiking was highly correlated. Together, these data indicate that tectal neurons that fire in sequence encode incremental dynamics in transforming spatiotemporal RFs.

So far, our data indicate that optic tectal neuron RFs can encode a range of stimulus features, from relatively simple ON or OFF stimuli to spatially and temporally complex stimuli, and that neurons which spike in a consistent sequence encode similar RF features. We investigated whether it is possible to build a logical diagram that generates spatiotemporal features of RF maps from the temporal sequences of firing. We considered two key aspects of our data in the model: one, neurons encode a variety of RF features; two, that spike firing sequence and timing are determinants in assembling spatiotemporal features of RFs. We schematized a raster plot to show the contribution of spike sequence to the logical diagram: neurons that consistently fire earlier or later than pairs of neurons with similar spike timing are candidates for respective upstream or downstream positions in divergent or convergent nodes (Fig. 4F, left: Spike Sequence data). We also assigned RF map features to neurons, denoted as A, B and C or combinations of these features (Fig. 4F, middle: RF Map data). We then organized the cells to form convergent and divergent nodes that generate a network with cumulative features of the RF maps (Fig. 4F, right: Logical diagram). Input from upstream divergent nodes that is shared between neurons with similar spike timing would increase the similarity of RF maps in those neurons (Fig. 4F, right: AB and AD). Similarly, convergent input from different upstream nodes would increase RF complexity (Fig. 4F: AB and ABC).

We evaluated this model using two independent strategies: in the first strategy we plotted RF similarity versus the relative number of shared earlier or later nodes between pairs of neurons (Fig. 4G). While RF similarity was correlated with increasing shared divergent nodes (Fig. 4G, top), RF similarity was not correlated with the shared convergent nodes (Fig. 4G, bottom). In particular, pairs of neurons that do not share an earlier node had almost no similarity in RFs (Fig. 4G, top, arrow). This analysis is consistent with the idea that the time-locked divergent node is a bifurcation point in information flow, as presented in the logical diagram.

In the second strategy to test the model, we asked whether candidate downstream neurons identified by consistent temporal firing sequences include different RF components seen in the individual upstream neurons, indicating that the downstream neurons may serve as a convergent node in the logical diagram (Fig 4F, right: AB and ABC). The Group 2 neuron pairs in Fig. 4E fire in a time-locked manner and have RFs with minimal similarity, raising the possibility that outputs from these neurons converge on downstream neurons (*26, 27*). We identified triads of neurons, in which a pair of neurons with similar spike timing both spiked with a consistent interval before the third neuron, a configuration consistent with a pair of candidate upstream neurons converging on a target (Fig. 4H, left, top panel). We compared the RF features in the candidate upstream and downstream neurons and applied the Moore-Penrose inverse matrix to identify the best linear representation of the RF features, A and B, that were represented in C (C vs C’ in Fig. 4H, left, bottom panel). This was compared with the consistency of spike timing between the candidate upstream pair of neurons and plotted in Fig 4H, right. The plot identifies a cluster of neurons in which firing consistency is significantly correlated with the contribution of upstream neurons to their later nodes’ RFs. The pairs of neurons that share common later nodes have less similar RFs (Fig. S6M), matching the feature of the Group 2 pairs in Fig 4E. Cells that do not have significant contributions from A or B may receive inputs from other nodes. These results show that subsets of neurons in the tectal circuit are highly time-locked and that the temporal coherence of firing predicts key nodes in a logical diagram based on RF properties. This suggests a model in which processing temporal information through circuits with time-locked sequential activity can integrate spatial and temporal information into complex spatiotemporal visual RFs (Fig. S7). The model circuit consists of repetitive motifs in which neurons share several common basic features of spatiotemporal response properties, specifically that dynamic responses transition from ON responses to OFF responses (Fig 1) and that inhibition controls the sequence of information flow through each motif (Fig 3). In addition, the motifs in the model circuit are organized in logical layers, which represent sequential encoding of spatiotemporal information.

### Plasticity of spatiotemporal filters

The data described above indicate that the timeline of input information is represented in the spike sequence in the circuit and that the sequence predicts the logical flow of information represented by the neurons. It is widely recognized that sensory experience alters neuronal response properties (*4, 28–32*). We sought to learn whether the temporal dynamics of spatiotemporal response properties might be tuned to predominant temporal features of environmental stimuli. Recordings of ON-OFF RF maps in response to unbiased reverse correlation analysis show that the ON-OFF RF maps are temporally dynamic. The peak of the OFF RF emerged around 100-200ms after stimulus onset with relatively little variance, while the peak of the ON RF showed significantly greater variance than the OFF response (Fig. 5A, B). We then tested whether the ON-OFF response interval is plastic in response to training with ON-OFF stimuli with different intervals (Fig 5C). The magnitude of the training-induced plasticity was highly correlated with the training stimulus (Fig 5D). Furthermore, the range in the training-induced changes in ON response times was larger than the range in the training-induced changes in timing of OFF responses, suggesting that plasticity in the timing of ON responses may underlie the tuning of temporal dynamics (Fig. 5E). Together, these data indicate that neurons encode the time scale of visual inputs and tune their temporal response properties to prevalent features of the environment.

**Fig. 5.**
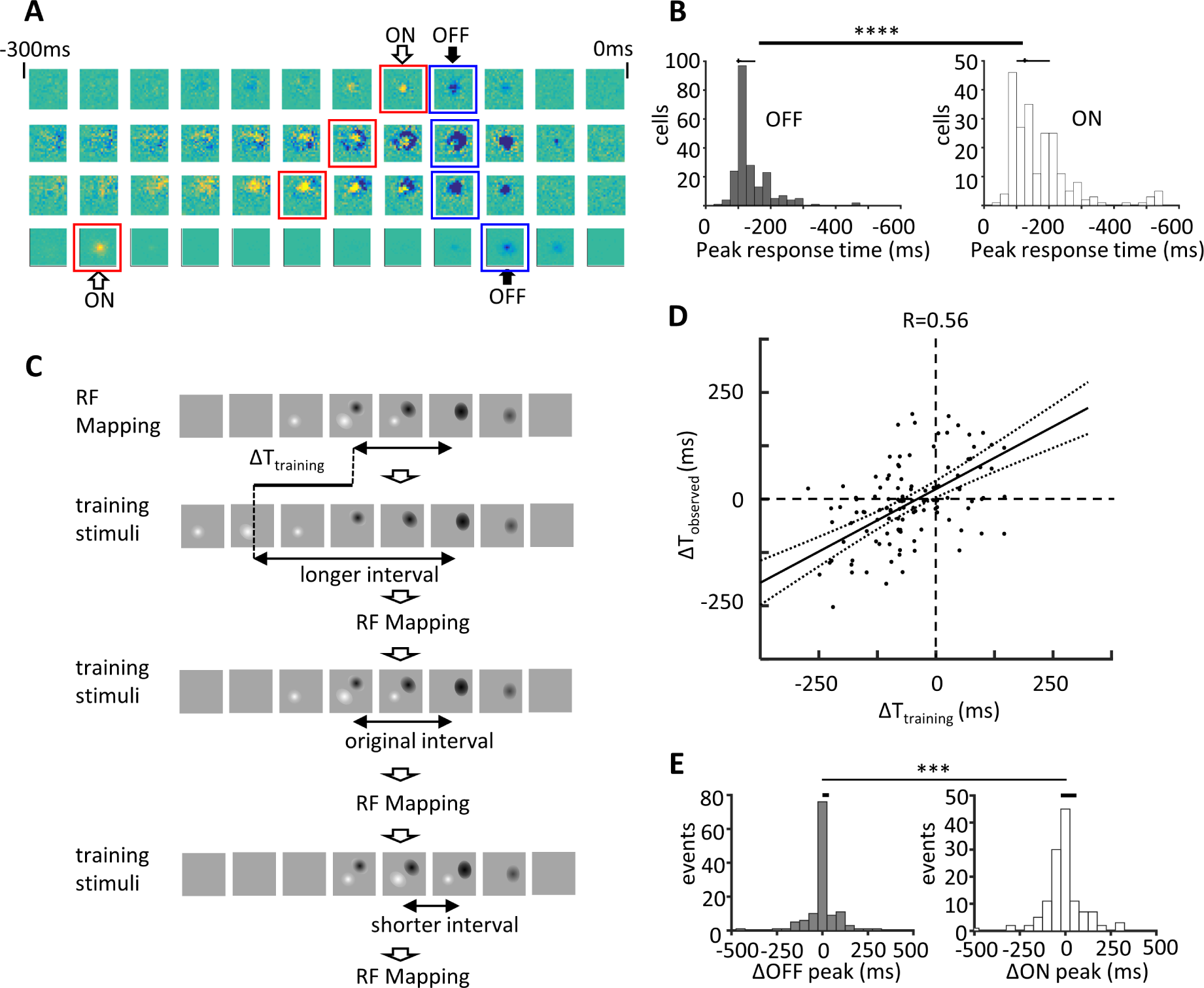
Sensory experience tunes the temporal features of visual responses. **A, B.** Temporal dynamics of OFF-ON RF transitions. **A.** Examples of ON-OFF spatiotemporal RF maps determined by unbiased reverse correlation showing a range of temporal intervals between the ON and OFF responses. **B.** Histograms of response times for OFF and ON stimuli for units with ON-OFF RFs. The variance of OFF responses is significantly less than the variance of ON responses. Interquartile range (IQR) = 50.0 ms (OFF), 100.0 ms (ON), P = 7.52e-14 (F-test), N=214 units in 38 animals. **C-E.** Visual training drives plasticity of the ON-OFF interval. **C**. Design of training stimuli. The original ON-OFF movie stimulus was determined using unbiased reverse correlation analysis from the spatiotemporal RFs (top row). To generate training stimuli with longer or shorter ON-OFF intervals, ON and OFF segments in the original movie images were shifted in time (≤T training). To generate the training movies, ≤T training stimuli were embedded within a ∼520ms image session so the OFF stimuli align in time across the movies with shorter or longer training intervals between the OFF and ON stimuli (top and bottom rows, ‘longer’ and ‘shorter’). Movies (700ms) were played repetitively with a 1.5 second gray pause after each movie stimulus. One session consists of four repetitions of the same movie stimulus followed by 15 seconds of gray. Each session is repeated 10 times. We presented the training stimulus for 2 h and mapped RFs after each training set. **D.** Plot of change in the ON-OFF interval (ΛT_observed_) versus the difference between the original ON-OFF interval and the ON-OFF training stimulus (ΛT_training_). The training-induced plasticity was highly correlated with the training interval (R=0. 56, P=6.6e-12). **E.** Variance in changes in response times induced by training for OFF and ON responses. IQR =25.0 ms (OFF), 75.0ms (ON), P=1.6e-4, F-test. N=19 x 3 trials, 19 animals.

## Discussion

Here we demonstrate that optic tectal neurons encode dynamic spatiotemporal image transformations, indicating that individual neurons in animals detect a variety of complex image trajectories by integrating inputs over extended time periods. Processing both spatial and temporal visual information enables the system to encode the trajectory and speed of moving objects. Many optic tectal neurons recognize spinning stimuli, and they demonstrate speed-dependent changes in the direction of the recognized stimulus, suggesting that the perceived aliasing of spinning objects may be neural circuit phenomenon. Based on our electrophysiology and calcium imaging data showing that tectal neurons fire in reproducible sequences to the preferred visual stimuli, that the temporal relationship of spikes between neurons matches the spatiotemporal features of RF maps across neurons and that inhibition plays a strong role in the generating the spatiotemporal specificity of the RF, we propose a circuit architecture in which nodes of interacting neurons compose a motif that is repeated both in parallel and sequentially through the circuit. This architecture enables encoding of stimuli with increasing spatiotemporal complexity as the number of nodes increases. Furthermore, temporal information is encoded by the ON-OFF response interval and is subject to experience-dependent plasticity, consistent with the idea that plasticity mechanisms strengthen stimulus motifs that are predominant in the environment.

Sensory circuits are thought to represent multiple dimensions of information organized in a hierarchical manner. For example, spatial information in visual cortex is represented in hypercolumns, where OFF RFs represent object locations, whereas OFF-ON vectors encode additional spatial information, such as object orientation (*2, 3, 33, 34*). Our study extends these principles of hierarchical organization of visual information to the temporal domain. OFF stimuli evoke spikes with a consistent latency following stimulus presentation, whereas the intervals between OFF and ON responses are variable. This suggests that the relatively stable OFF responses serve as an anchor for encoding temporal information, reminiscent of the idea that in the spatial domain, the OFF RF represents an anchor for the OFF-ON vector of object orientation (*3,34*). The plasticity experiments further support the idea of hierarchical organization of temporal information. Here, the timing of the OFF response was relatively stable and the timing of the ON responses changed following visual experience-dependent training, suggesting that OFF responses encode the timing of an event while the ON-OFF interval encodes duration of the dynamic stimulus. This analysis indicates that the presence of hierarchical relationships between different features of sensory information is an organizational principle with respect to both the composition and experience-dependent plasticity of information. Several concepts discovered in visual processing contributed to the recent remarkable progress in processing algorithms of static images, using deep learning (*35*). The coding rules found here show an example of a successful algorithm for detecting and encoding spatiotemporal information that may also be applicable to computer science.

## References and Notes

### Acknowledgments

We thank Dr. Carlos Aizenman, Dr. Edward Ruthazer and Dr. Nick Spitzer for comments on the manuscript and helpful discussions.

### Funding

National Institutes of Health EY011261 (HTC)

National Institutes of Health EY027437 (HTC)

National Institutes of Health EY031597 (HTC)

### Author contributions

Conceptualization: MH, HTC

Methodology: MH

Investigation: MH

Visualization: MH

Funding acquisition: HTC

Supervision: HTC

Writing – original draft: MH, HTC

Writing – review & editing: MH, HTC

### Data and materials availability

All data are available in the main text or the supplementary materials.

## Supplementary Materials for

### Materials and Methods

All experimental protocols were approved by the Institutional Animal Care and Use Committee at The Scripps Research Institute.

#### Animals

Xenopus laevis tadpoles were generated from the lab colony or purchased from Xenopus Express (Brooksville, Florida) and reared from stage 23 at 21°C with 12h dark/12h light cycle.

#### Recordings

Experiments were performed at room temperature (20-22°C). Animals were paralyzed in 2 mM pancuronium bromide (APExBIO, Cat#1612) for 1 min and then were mounted on an adjustable stage located at the center of the recording chamber containing extracellular saline, (mM): 115 NaCl, 2 KCl, 3 CaCl_2_, 3 MgCl_2_, 5 HEPES, 10 glucose, and 0.01 glycine, 0.01 pancuronium dibromide, pH 7.2, osmolarity 255 mOSM. Four-chamber glass-capillaries (Hilgenberg, Germany) were pulled on a Sutter P-97 puller, filled with external solution and tips were cut to generate a flat end of ∼15-20 μm opening, equivalent to the diameter of ∼1 cell body per channel. The glass pipette was set in a custom-made AgCl electrode holder. Isolation of each channel was confirmed by a multimeter before use. The electrodes were connected to patch clamp headstages by extension cables. Patch clamp amplifiers (Multiclamp 700A, 700B, Molecular Device) were used for signal amplification. The data were collected by DigiData 1440A (Molecular Device).

Spikes were sorted with the unsupervised sorting methods developed by Quiroga et al (*36*). We detected spikes with an amplitude threshold after frequency filtering (300-3000 Hz) and calculated the wavelet coefficients. Three coefficients that best separated different units were used for clustering. We used an initial set of recordings to map the RFs, from which we generated the template waveforms used for spike-sorting of data acquired subsequently in the same experiment. The clean glass pipette tips formed tight seals with the surface of neural tissues, which stabilized contact for several hours and excluded signals from cells outside of the contact area. 1-3 units were usually isolated from the four-channel electrodes, consistent with the tip sizes of the recording pipette.

For drug perfusion, external solution containing the reagents was added to a 60 ml sub-chamber connected to the recording chamber through two lines of tubes. The external solutions flowed into the recording chamber from the sub-chamber for 15 min and recordings were initiated after another 10 min.

#### Receptive Field Mapping

Spatial receptive fields were mapped using methods modified from previous studies (*16*). Images were generated on a computer and projected with a 3M MPro 110 miniprojector onto the back-projector screen on the recording chamber covering 90 degrees of the visual field. Tadpoles were stabilized on a platform positioned 15 mm away from the screen. Image brightness was controlled by a neutral density filter (0.07-0.33 mW/cm^2^). Timing of the image presentation was detected with a photodiode positioned at the corner of the screen. For receptive field mapping at coarse resolution (Fig. S2), white squares (viewing angle: 11.25°, 0.325 mW/cm^2^, background: 0.128 mW/cm^2^) were presented in random 8×8 grids for 1.5 sec with a 5 sec interval for 64 times without duplication. Spike numbers from three repetitions were averaged. For mapping at higher spatiotemporal resolution (Fig. 1-4), we used an unbiased sparse noise visual stimulation protocol, presenting a random series of stimuli composed of two white or black squares on a grey background (viewing angle: 9°, white: 0.325 mW/cm^2^, black: 0.069 mW/cm^2^, background: 0.128 mW/cm^2^) at random in 20×20 grids for 400-600 msec for 4000-16000 times (*3*). The centers of the two squares were separated by 13.5 to 22.5° of the viewing angle. Images of stimuli were interleaved with 400-1000ms intervals to detect extended temporal response dynamics. RF maps were calculated by reverse correlation (*3*, *17*) and presented as a spike triggered average where the image in each time window is the average of stimuli that evoked spikes at 0 ms (*18*).

#### Analysis of direction selectivity

To determine the location in the visual field to present stimuli, the gravity centers of ON and OFF RFs were identified from the reverse correlation analysis of the sparse noise stimuli. A circle of grating was presented so that the darkest and the brightest peaks of the grating matched the gravity centers of ON and OFF RFs, respectively. The diameter of the circle was twice the distance between the gravity centers. After presenting a still image of the grating for 5 sec, the grating drifted for four cycles. Total spikes during the movie presentation were summed and displayed in polar plots (Fig. S3).

#### Analysis of rotation direction selectivity

Movies of a rotating sphere were presented as stimuli. The sphere was generated by fitting (x, y, intensity) data of RF map pixels from the reverse correlation analysis of the sparse noise stimuli. The sphere was painted with a white to black gradient from one pole to the other pole. The sphere was rotated on a gray background (white: 0.325 mW/cm^2^, black: 0.069 mW/cm^2^, background: 0.128 mW/cm^2^). The frame of the RF map at the middle of ON and OFF response peaks was selected to determine the peaks in the black and white grating. The first frame image was presented for 5 sec before the start of rotation. 10 cycles of movies were presented. Total spikes evoked by the movie were counted. Stimuli with different speeds were presented in a pseudorandom order. Each set was repeated eight times and compared with the response to the opposite rotation.

#### Generation of a series of movie stimuli in a principal coordinate

Spatiotemporal spike-triggered average (STA) in response to the sparse noise stimuli was used to generate the basic stimuli. The background level was set at 0.128 mW/cm^2^. The brightness was scaled so that the brightest and darkest pixels in the stimuli were 0.325 or 0.069 mW/cm^2^, respectively. A sequence of receptive field maps was transformed into a trajectory of principal components. The eigenvectors of the coordinates were generated from vectors generated by linearizing all images in the spatiotemporal RF Maps. To generate a set of trajectories by rotation (0, 45, 90, …315 degrees), a center time frame (t_center) was defined as the middle of the peaks of OFF and ON responses (defined as t_off and t_on, respectively). Position (P [t_center]) did not move in the rotation. Other positions (P [t]) rotated on a circle whose diameter was a line that connected each position and the opposite position on the trajectory (P [2 x t_center – t]). The line connecting P[t_center] and center of P[t_off] and P[t_on] was defined as the normal vector of the circle for the rotation. Therefore, 180-degree rotation generates the reverse sequence.

#### Simulation of evoked responses

To predict responses (Fig. S4B), the cross-correlation between the optimal movie stimulus and the presented spinning stimulus was calculated. The response duration to calculate the optimal movie stimulus was determined by applying Otsu’s method to the peri-stimulus time histogram (PSTH) during 0-600ms from the random noise RF mapping. The maximum cross-correlation of the optimal movie stimulus and the spinning movie were calculated to predict the response profiles.

#### Training by image sequence stimuli

Spatiotemporal RF maps, which last 775 ms, were obtained by presenting the sparse noise stimulus used in Fig. 1. OFF and ON components of RF maps were separated and the OFF-ON interval was shifted to longer or shorter intervals. The image sequences were merged to generate the training stimuli of longer (+75 ms), original, and shorter (-75 ms) OFF-ON intervals. One movie stimulus was presented after 100 ms background and was followed by background for a total of 1.5 s for the movie. The movie was presented four times every15 sec. This set continued for 2 hr for each training movie sequence. Spatiotemporal RFs were mapped after every 2 hr training session. Training sessions that included movie stimuli in which the OFF and ON stimuli were presented in a reversed sequence were excluded from the regression analysis.

#### Calcium imaging

To transfect tectal neurons for calcium imaging experiments, stage 45 tadpoles were anesthetized in 0.02% MS-222 (Tricane metanesulfocate; Sigma-Aldrich, St. Louis, MO) and electroporated with DNA plasmids (pCMV::GAL4: 2µg/µl, pUAS::GCaMP6f: 2 µg/µl (*37*), pCMV::turbo RFP (HC1975) 2 μM/ul). Electroporation of the optic lobe was accomplished using unipolar current pulses, as described (*38*). Animals were raised in 12 hr light 12 hr dark cycles until stage 47. The animals were paralyzed in 2 mM Pancuronium Dibromide in Steinberg’s solution for 1 min and were mounted on an adjustable stage located at the center at the imaging chamber containing extracellular saline.

Pairs of image stimuli were projected on a back-projecting screen on one side of the chamber, as described for the electrophysiological recordings. To avoid interference with the PMT, a band pass filter (607/45 nm) was set on the projector (3M MPro110) with a focus adjust lens (f=150 mm). GCaMP6f fluorescence images were collected at 30 Hz on a two-photon microscope (Scientifica) with image acquisition software (Vidrio Technologies, ScanImage5). A 940 nm laser (Coherent) was used for excitation.

To digitize Ca^++^ signals, image frames were registered by 2D-cross correlation. An image representing a standard deviation along t-axis was generated from a XYT stack. To identify regions of interest (ROIs), contour lines at various values (5% step) were made on the image and non-overlapping ROIs were selected manually. ROIs of neuronal cell bodies or thick primary dendrites that showed less than 10% coactivity with other ROIs were selected for analysis to avoid potential duplicated measurement from the same neurons. To identify Ca^++^ spikes, frames that met the following criteria were identified. The intensity change between neighboring image frames was calculated as dF. Considering a sequence of 3 frames, 1. The increment of the signal from the previous frame was mean+3 standard deviations or larger. 2. The signal from the frame immediately previous to the candidate spiking frame was larger than its previous frame. 3. Signals subsequent to the spiking frame were larger than the spiking frame for at least 300 ms. This excludes Ca^++^ transients with a gradual increase or plateau. Amplitudes of the signal were normalized to the average baseline fluorescence over -5 s to + 5s, relative to the Ca^++^ transient. This normalization was robust against transient perturbation of baseline compared with the normalization by F0 (Fig. S6A, B). To calculate the spatial RFs, each stimulus image was multiplied by the amplitude of Ca^++^ spikes and summed. The color values in Figs. 4C, S6I were calculated as follows. The typical firing sequence between each pair of cells was determined by the average relative spike timing. One spike was selected and other cells that spiked within ±100 ms were identified. We calculated the fraction of spikes within ±100 ms that follow the typical firing sequence as a matching index. We used a bootstrap test to unify the statistical analysis within a dataset, in which the percentages of significant difference between subsets of data were compared. ICA analysis (Fig 4E) was performed using FastICA (research.ics.aalto.fi). The origin was defined to minimize the residual.

---

**Fig. S1.**
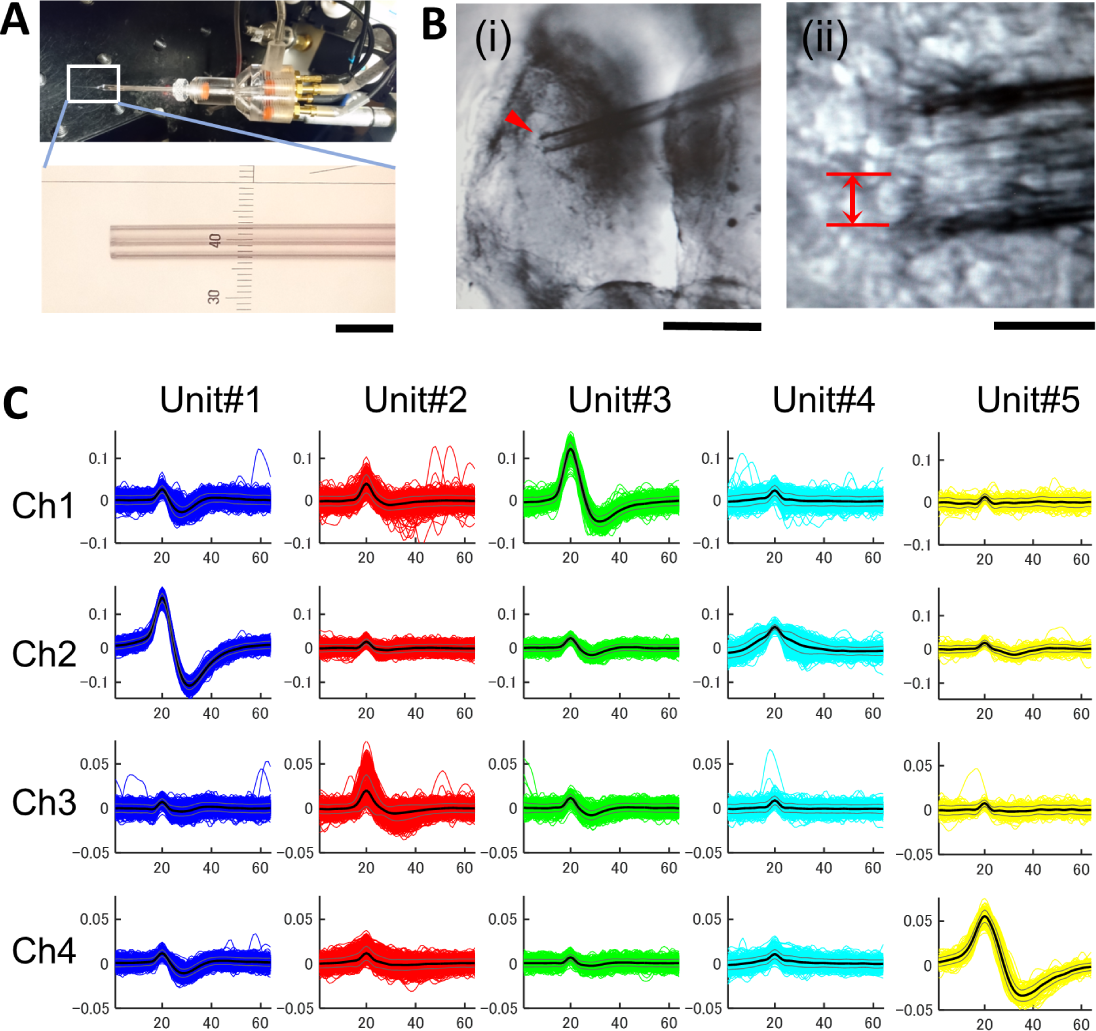
Visual receptive field mapping in optic tectal neurons in Xenopus tadpoles. **A, B.** Images of a juxtacellular four-channel glass electrode (A) positioned within the optic tectum (B-i, arrowhead). The size of a neuronal cell body at the pipette tip (B-ii, red lines) **C.** Examples of sorted waveforms from juxtacellular multi-channel recordings using multi-channel glass electrodes. We typically detected ∼1 unit per channel which reduces uncertainty in spike sorting. Scale bar: 100μm (A), 200μm (Bi), 20μm (Bii). Modified from (*15*).

**Fig. S2.**
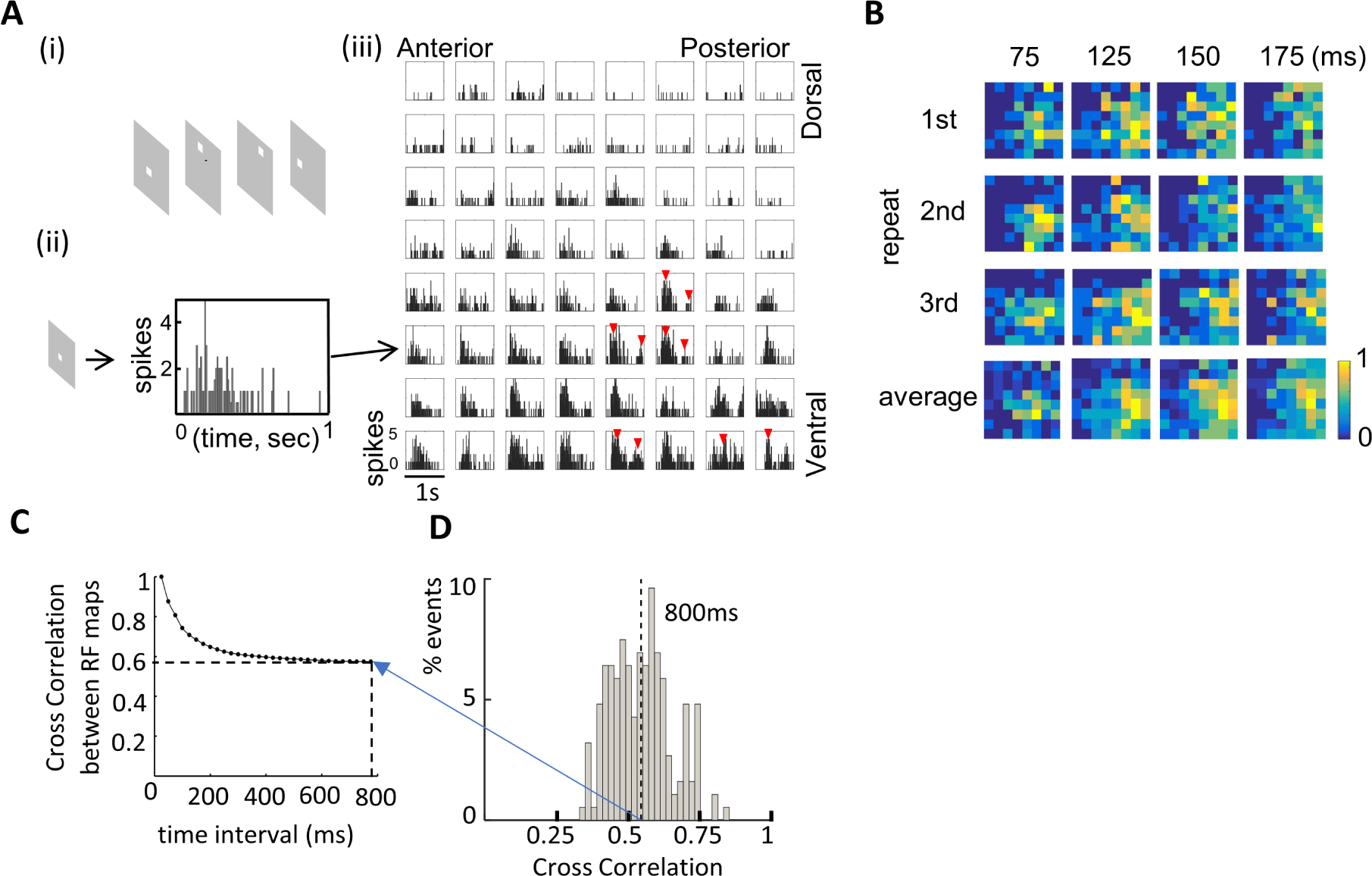
Spatiotemporal RF maps are variable over time. **A.** Classical method of mapping spatiotemporal RFs over 1 second epochs using cell attached patch recordings. (i) Small white square stimuli are presented in a matrix of pseudorandom locations in the visual field for 1.5 sec with a 5 sec interstimulus interval. (ii) Poststimulus time histogram of spike responses to a single stimulus over a 1 second recording period. (iii) Histograms of spike numbers over time for stimulus locations across the visual field. The matrix shows the temporal sequence of responses over 1 second of recording for stimuli presented in anterior to posterior locations in the visual field (left to right in matrix) and dorsal to ventral positions in the visual field (top to bottom in matrix). Red arrows identify multiple peaks of spikes across time in particular RF locations. **B**. RF Maps from three repetitions of the same set of stimuli and their average. Pseudocolor scale: spikes normalized to the maximum value (yellow) for each time bin. **C, D.** Quantification of changes in RF stimuli over time by cross correlation. 2D cross correlation between all combinations of RF maps across 25ms time windows were calculated for each unit and pooled (**C**, N=64 units). The histogram (**D**) shows the average cross correlation when 25ms time windows are sampled over 0-800ms after stimulus presentation. When the time range is 0-800ms, average is 0.54 (dotted line).

**Fig. S3.**
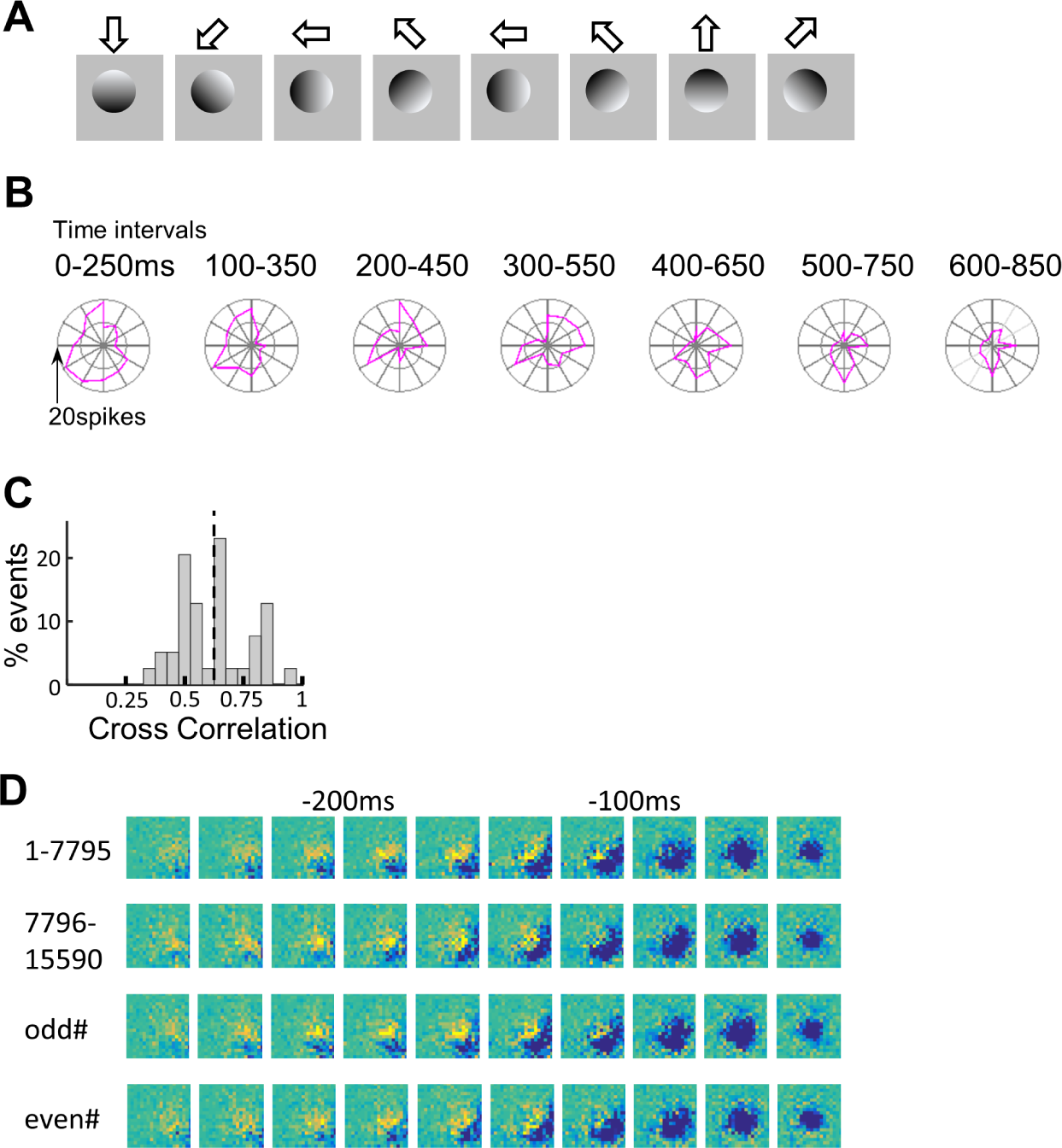
Direction selective responses are temporally dynamic. **A.** Moving 1 Hz gratings moving in different directions over 1 second were used to map direction selective responses. **B.** Maps of direction selective responses over time were generated from data collected over 250 ms windows shifted by 100 ms from left to right. To define the location and spatial frequency of the test stimulus, gravity centers of ON RFs and OFF RFs were identified from the RF map frames in which ON and OFF responses were highest, respectively. The stimulus grating image was made so that the line connecting ON and OFF gravity centers matches one cycle of the grating (53.5 deg of the visual field). **C.** Histogram of the average normalized cross-correlation of tuning profiles for 1 Hz dataset (vector of spike numbers for individual angles) across time windows, using the time window with the highest orientation tuning index as the reference (n=39 units). Units with >9 spikes for all angles were included. Dotted line: mean (0.63), Standard deviation = 0.15. **D.** RF maps generated from subsets of stimulus sessions. A total of 15590 sparse noise images were presented. RF maps generated from the stimulus sessions 1-7795 and 7796-15590 are comparable, and RF maps generated from even and odd numbers of the stimulus sessions were comparable.

**Fig. S4.**
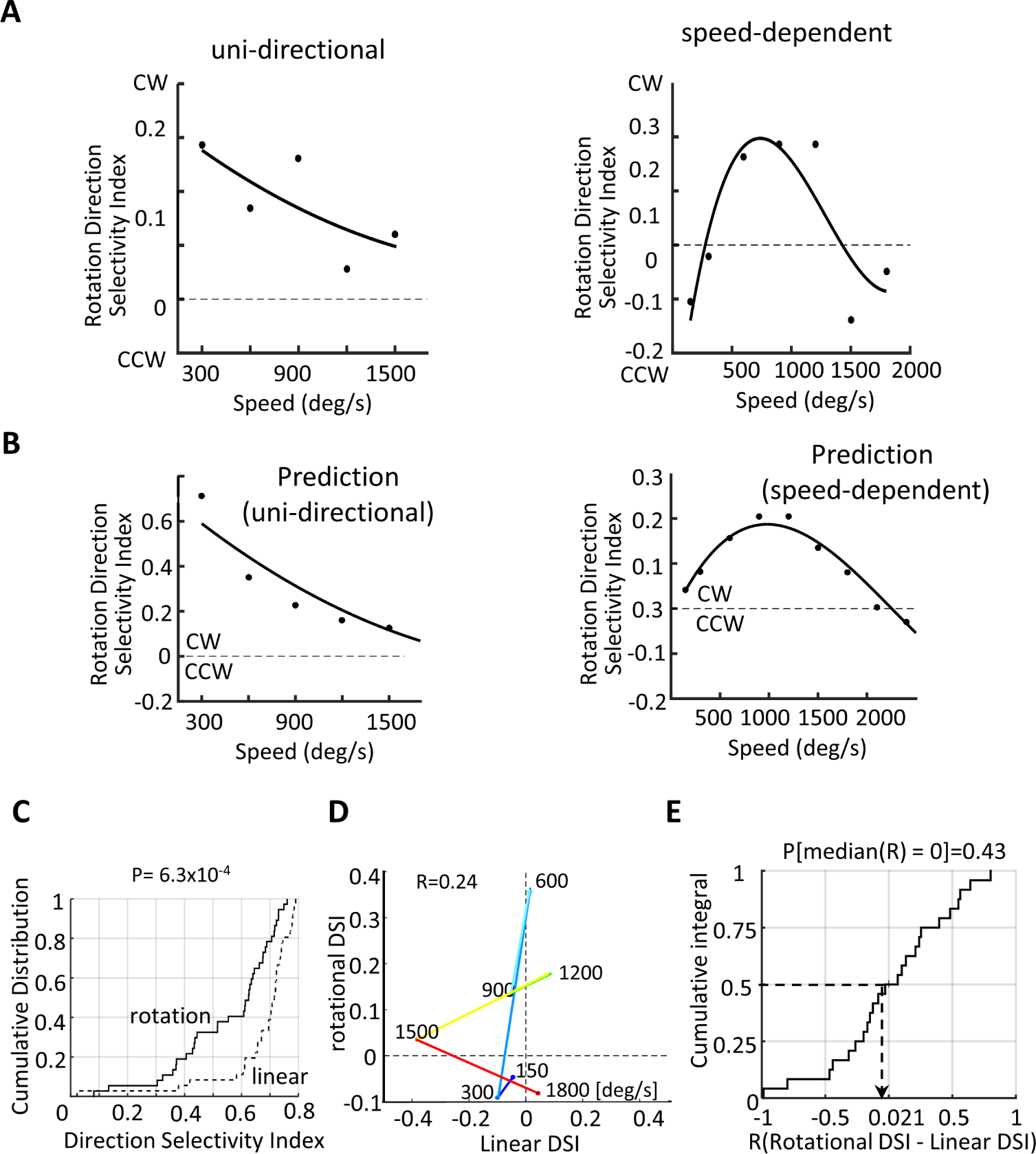
Detailed analysis of rotation preference. **A, B.** Rotation direction selectivity index calculated from Fig. 2C **(left panels)** and predictions from spatiotemporal RF maps **(right panels)**. Examples of neurons are shown in which the preferred rotation direction of the stimulus is consistent (A) or switched between clockwise and counterclockwise depending on rotation speeds (B). Rotation Direction Selective Index (RDSI) is calculated as follows: RDSI = (nSpikes(CW) - nSpikes(CCW))/ (nSpikes(CW) + nSpikes(CCW)), where nSpikes(CW) and nSpikes(CCW) are the numbers of spikes evoked by clockwise and counterclockwise rotating image stimuli, respectively. Prediction of the rotational direction selectivity index was made from cross-correlation between the rotating stimuli. **C.** Cumulative distribution of absolute DSI of responses to rotation or linear motion (P<0.01, Komogorov-Smirnov test, N=24). **D.** Plot of RDSI compared to linear DSI at different speeds, for the neuron analyzed in Fig 2D. No correlation was observed (R=0.24). **E.** Population analysis of the cumulative frequency of correlation coefficients (R) between rotation DSI and linear DSI, showing the random distribution of the correlation coefficients. The median value =0.021 is not significantly different from 0 (p=0.43) (bootstrap test, 10^4^ repeats).

**Fig. S5.**
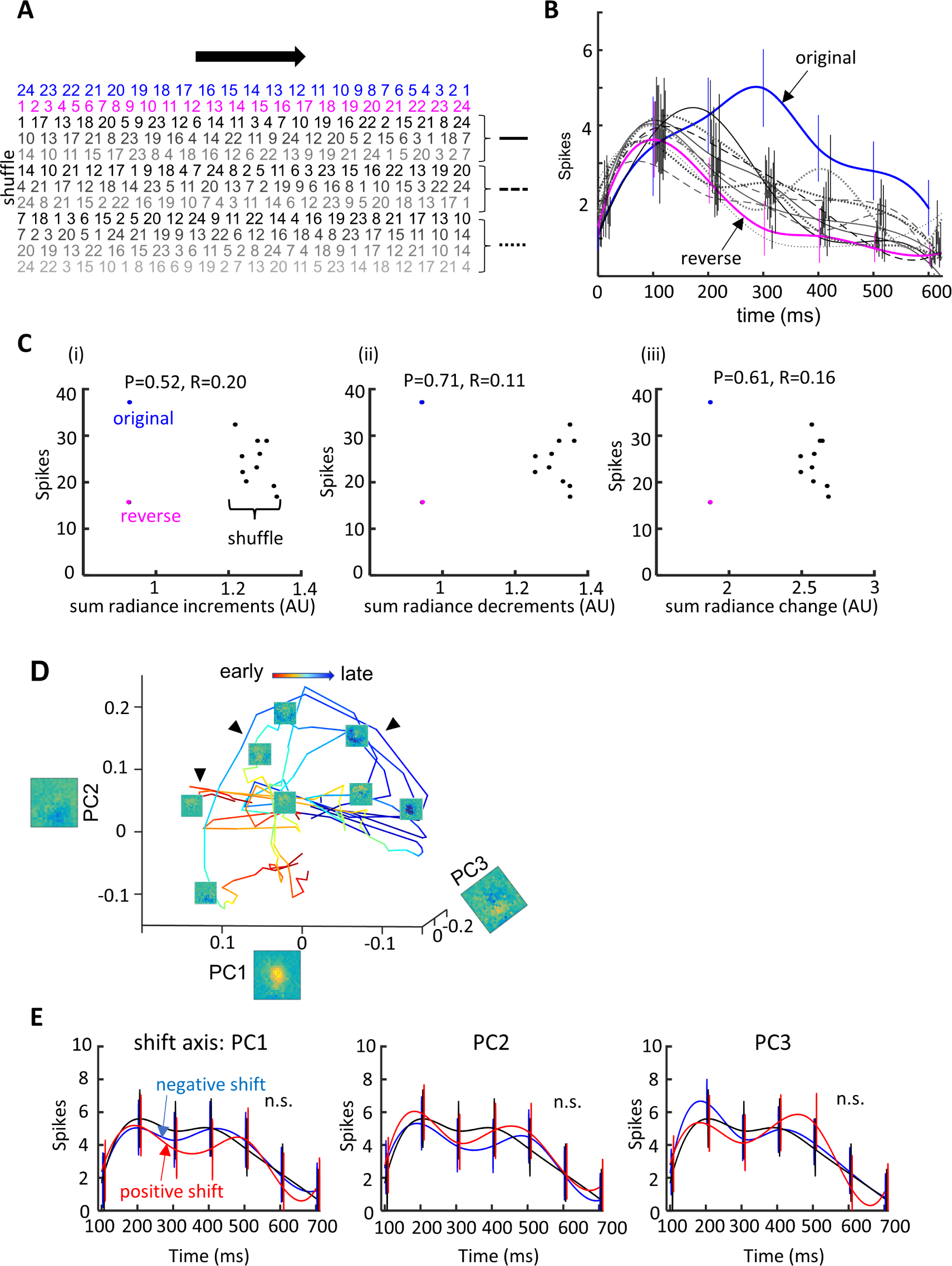
Tectal neurons recognize sequence specific image transitions. **A, B.** Responses to original (blue), reverse (magenta) and the full range of shuffled (black-gray) frames of movie stimuli for spatiotemporal RF maps. **A**. Sequences of frame numbers for original (green), reverse (red) and shuffled (black) frames. Shuffled sequences are bracketed into 3 groups, shown on the right, and individual sets of shuffled stimuli, shown in gray scale, are plotted in B. **B**. Plot of average spike numbers per 100 ms time window over the 600 ms movie stimulation period for the original (blue), reverse (magenta) and the shuffled (black-grey) movies. Bars show standard deviation of the mean. **C.** Changes in radiance do not account for changes in responses to forward, reverse or shuffled stimulus sequences. Plots of total spike numbers across total radiance changes (i: increments, ii: decrements, iii: absolute changes) and for the original (blue), reverse (magenta) and shuffled (black) image sequences. AU=arbitrary units. Data from Fig. 3B. No correlation was observed between the total radiance change and the total spike numbers (P (R=0) and R values in each panel). In general, total changes in radiance in the original (blue data points) and reverse movies (magenta data points) were smaller than the shuffled movies (black data points). The largest responses were evoked by the original movie and the smallest responses were evoked by the reverse movie, as shown in Fig. 3B, C, indicating that radiance change does not explain the difference in the responses. **D.** Sequence of transitions of principal coordinates of spatiotemporal response properties (PC1:17.0%, PC2: 9.6%, PC3: 5.7%) indicates that the spatial patterns vary over time (from red to blue, early to late). One segment is 25ms. All cells share a similar trajectory during earlier (red) and later (blue) timepoints but diverged at ∼100-200ms and tracked different pathways. Subsets of paths were shared by several trajectories (arrowheads). The images on the axes (PC1-3) show the principal component in the RF maps. Data from a subset of 7 out of 54 units were plotted to avoid dense overlay. **E.** Parallel shifts of stimulus movies in the principal coordinate did not change evoked responses, graphed as spike numbers. Responses to the original movie stimulus (black) and stimuli based on positive and negative shifts in principle components (red and black lines, respectively). The modified movies were made by shifting the trajectories two times the range of each trajectory (Fig. 3F) in each axis of principal components. The amplitude of the shift was equal to the dynamic range of the trajectories in each axis. Data are the average spike number from three repeats of each stimulus. X axis is time after movie started. The numbers of the spikes did not change significantly in response to parallel shifts of the trajectories. P=0.99 (PC1), 0.95 (PC2), 0.98 (PC3), ANOVA. N=8 neurons from 8 animals.

**Fig. S6.**
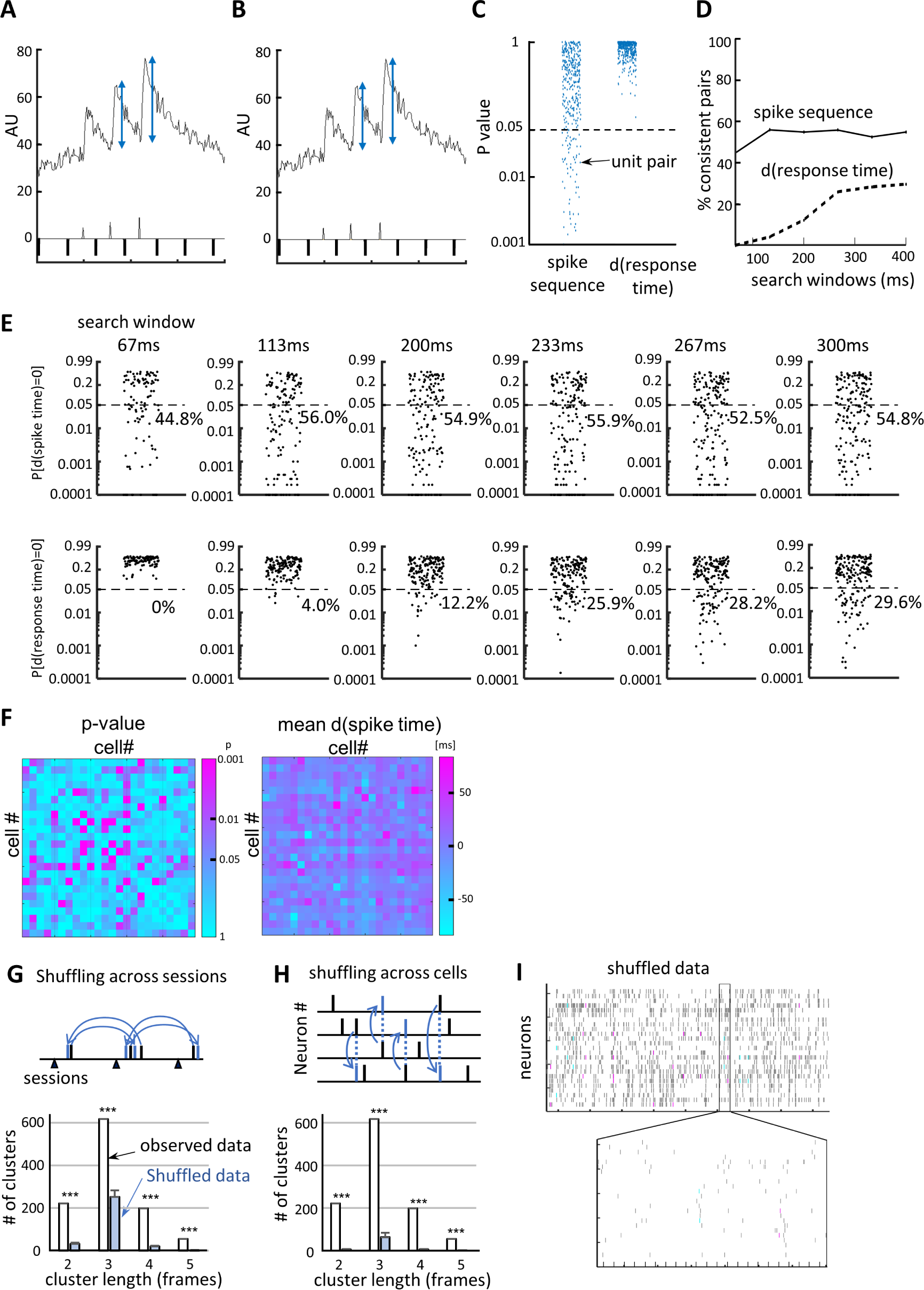
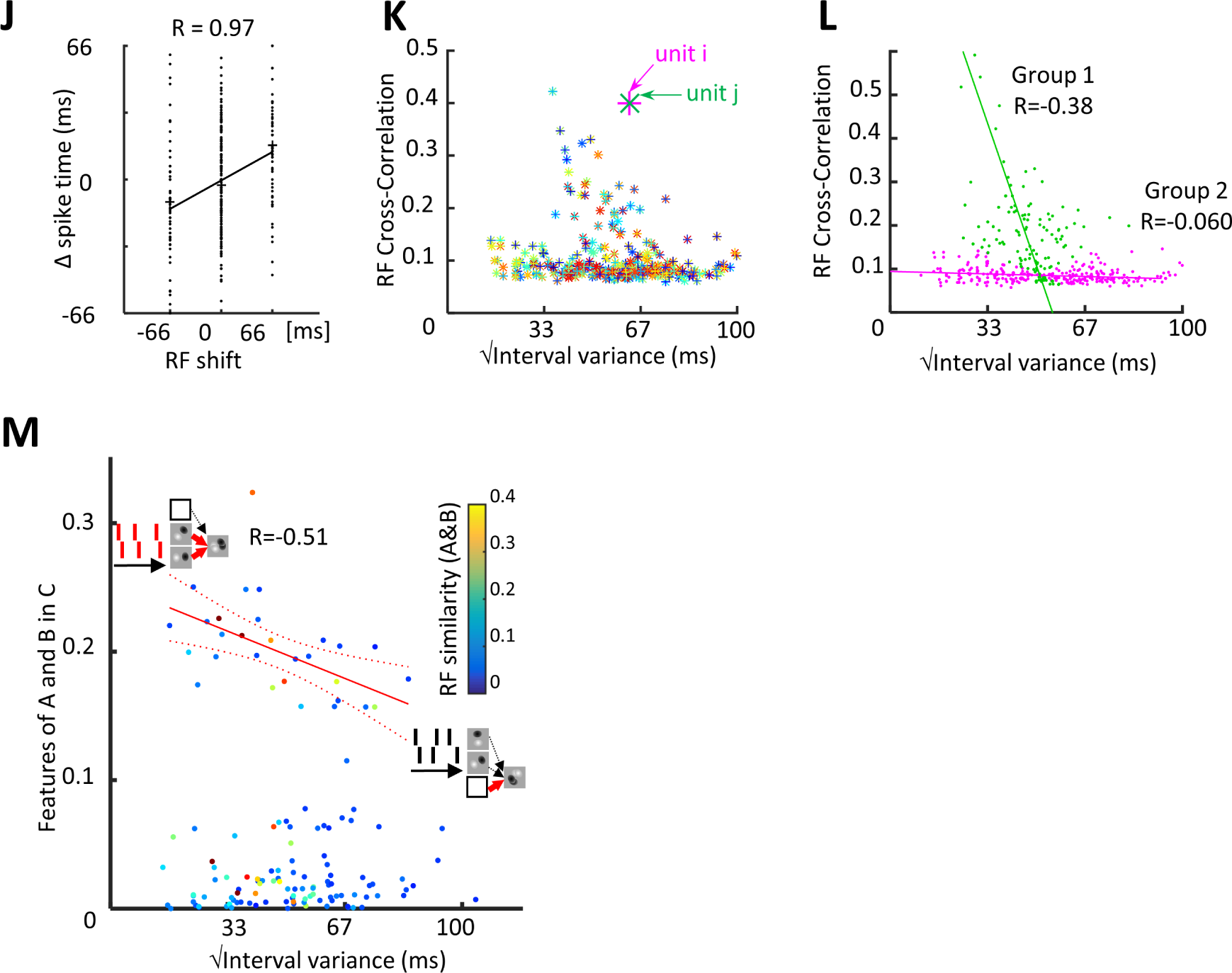
Analysis of calcium imaging Data. **A. B.** Comparison of two methods for background normalization used for spike detection: Normalizing signals in each frame by the rolling average fluorescence of images over 10 sec, with 5 sec before and after each frame (**A**) or by F_0_ (**B**), where F_0_ is the fluorescence in the previous frame. Increased amplitudes of spikes were reliably detected when data were normalized to a 10 second rolling average of fluorescence (inset along X axis). This avoids potential influence of the residual Ca^++^ signal from a previous spike on the F0 value used for normalization. Downward spikes show stimuli presented every second. **C**. Pairs of neurons fire in a reproducible sequential order. Spiking activity from pairs of neurons were searched within 100ms window. Left: Each data point indicates that the probability that the pair fires in a consistent sequence. Pairs with p< 0.05 are considered to fire in a reproducible sequential order. 22.0% of pairs fired in reproducible sequential order. Right: To evaluate the possibility that the appearance of sequential firing between neurons arises from consistent differences in the response times after stimulation, we tested the probability that stimulus-response times were different between neurons in each pair. Each data point indicates that the probability that stimulus-response times were different between neurons in each pair. 0% of pairs exhibited a significant difference in average response time. (427 pairs from 4 animals). **D, E.** Extending the search window from 67ms to 400ms increases the detection of neuron pairs that fire in a sequence, but also increases the pairs of neurons in which the individual neurons have different response times after stimulation. **E.** Raw data for D. **F.** Matrices showing relative spike timing across neurons over 100ms (left) and differences in response latencies between the individual neurons in each pair (right). Right: matrix of p-values evaluating consistency of the order P(dt=0) calculated by bootstrap test (10^4^ repeats). 28.7% of pairs spiked with consistent sequential order (P<0.05), while only 0.22% of pairs showed significant differences in the latency from stimulus to spike time (56 cells from 4 animals) **G.** (top) Shuffling strategy. Response times after image presentation (black arrowheads) were shuffled across sessions. To test modest shuffling, the number of spikes in each session was unchanged. (bottom). Histogram of cluster lengths showed a significance of difference between the observed data and shuffled data (*** P <10^-5^; bootstrap test, 10^5^ repeats). Cluster length is the number of contiguous frames with consistent sequential firing. **H.** An alternative shuffling method. Here timestamps of spikes were randomized across cells (left). The number of trains the shuffled data was significantly smaller than the experimental data. (** P<10^-5^, bootstrap test, 10^5^ repeats). **I.** Raster of shuffled data using modest shuffling strategy in G. Few cell pairs (4.9±3.6×10^-4^ %, bootstrap test, 1000 repeats) fire in a reproducible sequential order (shown as very few colored datapoints). **J.** Plot of temporal RF shift and mean difference in spike timing between all pairs of neurons. N=4 animals. **K**. Plot of Fig. 5F showing identity of units in each pair by colors. **L.** Pooled data for Fig. 5F (N=4 animals).

**Fig S7.**
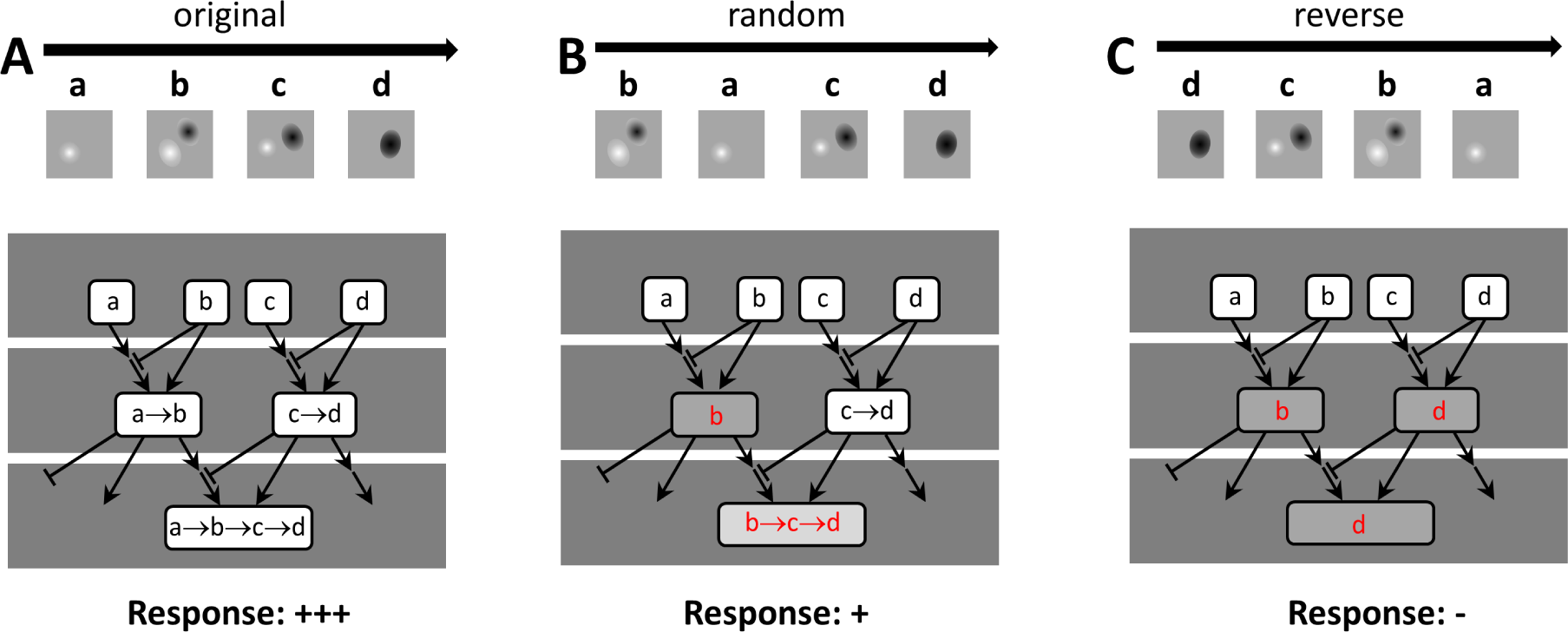
A schematic circuit model that encodes image sequences. A model consisting of repetitive circuit motifs, which include neurons or neuronal assemblies (white rectangles with letters) that represent receptive field features they encode in the stimuli, designated as a, b, c, d. Some neurons are excitatory (with arrows) and some are inhibitory (with bars). Motifs include nodes, sites of information processing, at which time-locked upstream inputs converge or from which time-locked downstream inputs diverge (Fig 5). Components of image sequences are encoded by upstream neurons (top rows). Convergent nodes (middle and bottom rows) generate responses that capture the temporal relation of responses in the upstream neurons. Inhibition is present upstream of convergent nodes and controls the sequence of response features encoded by the downstream neurons, consistent with the picrotoxin sensitive sequence selectivity observed in Fig. 3. A neuron that encodes the temporal sequence of “a→b→c→d” is located in the bottom row of panel **A**. The original image sequence activated the neurons in the top row in a temporal sequence, a→b→c→d (A). This sequence activates the neurons in the second and the last layers which encode more complex RF properties through a combination of convergence and delayed inhibition, so the complete image sequence is encoded. An image sequence with partial inversion (**B,** b→a→c→d) induces inhibition before integration of “a” and “b” in the second layer. This blocks input “a” from contributing to the activity of the left neuron in the second layer, so only input b, in red, in encoded. This results in the reduced response, b→c→d, red, in the bottom layer, which fails to encode the input image sequence. A reverse image sequence (**C,** d→c→b→a) induces inhibition at all convergent nodes and the minimal activity “d” is encoded by the neuron in the bottom layer. This schematic is sufficient to model responses to image sequences in which “a→b→c→d” (original sequence, **A**) and “d→c→b→a” (reverse sequence, **C**) evoke the strongest and the weakest responses, respectively, as recorded experimentally (Fig. 3B). More generally, the model explains how the integration of spatial information over time can occur through convergent and divergent nodes.

